# Spatial transcriptomics reveals injury-responsive compartments and coordinated immune–fibrotic signaling in ANCA-associated renal vasculitis

**DOI:** 10.1101/2025.09.19.677268

**Authors:** Yucheng Tang, Chunhua Zhu, Shuang Chen, Huimei Chen, Hai Qian, Aihua Zhang, Enrico Petretto

**Author notes:** Yucheng Tang and Chunhua Zhu contributed equally to this work. **Address for correspondence** Enrico Petretto, Institute for Big Data and Artificial Intelligence in Medicine, School of Science, China Pharmaceutical University, Nanjing 210009, China; Programme in Cardiovascular and Metabolic Disorders, Duke-NUS Medical School, Singapore, Singapore; Centre for Computational Biology, Duke-NUS Medical School, Singapore, Singapore **Email:**; Aihua Zhang, Department of Nephrology, Children’s Hospital of Nanjing Medical University, Nanjing 210008, China **Email:**.

## Abstract

ANCA-associated vasculitis (AAV) often presents with rapidly progressive glomerulonephritis, yet how immune and stromal programmes are organised within kidney tissue remains unclear. We applied spatial transcriptomics to renal cortex biopsies from patients with AAV of varying disease severity and normal-histology controls, generating a spatial map that localizes disease programmes *in situ*. Four disease-responsive compartments—immune/interstitial fibroblasts (IM/Fib), glomeruli, myofibroblasts, and vascular compartments—showed distinct compartment-specific signatures that correlated with histopathology at the patient level. We also identify coordinated immune–fibrotic signaling linked to severity. Within IM/Fib, the CXCR4–CD74 receptor complex co-localised with IgM⁺ cells, and lumican (LUM) co-localised with collagen I/III; both were validated by immunofluorescence and associated with fibrotic injury. These tissue-anchored signatures constitute candidate diagnostic/prognostic biomarkers, and show the utility of this foundational spatial transcriptomics map to generate testable hypotheses which will guide validation in larger, stratified, treatment-annotated cohorts.

## Introduction

Anti-neutrophil cytoplasmic antibodies (ANCA)-associated vasculitis (AAV) represents a group of rare systemic autoimmune diseases, characterised by inflammation of the small-to-medium-sized blood vessels^1^. Renal involvement occurs in >75% of patients, predominantly manifesting as rapidly progressive glomerulonephritis (RPGN)^2^. In pediatric AAV, renal involvement is more severe, with approximately 20%-35% of children progressing to renal failure, while another 20%-30% develop chronic kidney disease (CKD)^3^. However, children exhibit a higher capacity for renal function recovery compared to adults, which is potentially attributable to the greater plasticity of the immune system and the potential for earlier therapeutic intervention^4^.

A central immunological mechanism of AAV involves the breakdown of immune tolerance to neutrophil granule proteins, primarily proteinase 3 (PR3) or myeloperoxidase (MPO), resulting in the production of ANCA^5^. These ANCAs activate neutrophils, promoting vascular injury through the formation of neutrophil extracellular traps (NETs), oxidative burst, and degranulation^6^. Activated neutrophils also release autoantigens into the extracellular space, where they are captured by dendritic cells (DCs) and presented to naïve T cells^7^. This leads to the expansion of effector T cells, which infiltrate renal tissue and contribute to crescent formation, interstitial fibrosis, and tubular atrophy^8^.

Transcriptomic (single-cell and spatial) studies in AAV have identified key immune cell populations and provided insights into disease mechanisms. Single-cell analyses have highlighted disease-associated monocyte subsets (*FCGR3A⁺* and *FCGR1A⁺*)^9^, *SPP1⁺* lipid-associated macrophages that are associated with inflammation and fibrosis^10^, and cytotoxic T cells exhibiting pathogenic profiles in renal tissue^11,12^. Activation of the alternative complement pathway has been demonstrated in AAV^13^. Collectively, prior studies primarily focused on circulating immune cells, leaving unresolved how immune and stromal programmes are organised within kidney tissue and how this spatial organisation relates to injury severity and repair in AAV.

Current AAV treatments rely on immunosuppressive therapy, typically cyclophosphamide with high-dose glucocorticoids, which improves survival but increases infection and malignancy risk^14,15^. The use of the anti-CD20 monoclonal antibody, rituximab (RTX), yields comparable renal outcomes, with more effective B-cell depletion and better ANCA seroconversion^16^. Avacopan is an oral C5a receptor antagonist that effectively blocks the activation of neutrophils and the release of NETs, thereby reducing inflammation and tissue damage in AAV^17^. It was authorised for adjunct use with cyclophosphamide or rituximab, combined with a glucocorticoid dosage lower than standard protocols. Yet 10–30% of patients fail to achieve remission, remain glucocorticoid dependent, or progress despite therapy^18^, and multiple immune-targeted agents are still in development^12,19,20^. Since current therapies improve survival yet leave a substantial fraction of patients with persistent activity, glucocorticoid dependence, and progression to CKD, there is a critical need for tissue-level biomarkers that quantify severity and resolve immune–fibrotic signaling *in situ*. We therefore hypothesized that spatially resolved transcriptomic signatures within the renal parenchyma would reveal physiologically relevant biomarkers and pathogenic circuits not detectable in blood. To test this hypothesis, we profiled rare, biopsy-limited renal cortex specimens from patients with AAV using 10x Genomics Visium, defined AAV-responsive regions and constituent cell types, and linked regional programmes to histopathological severity. We identified cell-type-specific markers linked to disease pathology, and prioritized ligand–receptor interactions that serve as biomarkers of disease severity in renal biopsies and as potential therapeutic targets in ANCA-associated vasculitis.

## Results

### Spatial transcriptomic profiling of kidney biopsies in ANCA-associated vasculitis

We categorized AAV patients into mild (*n* = 2) and severe (*n* = 3) groups and analysed kidney cortex biopsies alongside those of controls (*n* = 4) (Supplementary Table 1). The controls, obtained from patients with hematuria, showed normal renal histology without sclerosis or atrophy. In contrast, mild AAV biopsies displayed segmental sclerosis and tubular atrophy, while severe AAV consistently exhibited glomerular crescents, reflecting severity-dependent tissue damage (Fig. 1a). None of the patients had received treatment at the time of biopsy collection.

**Fig. 1:**
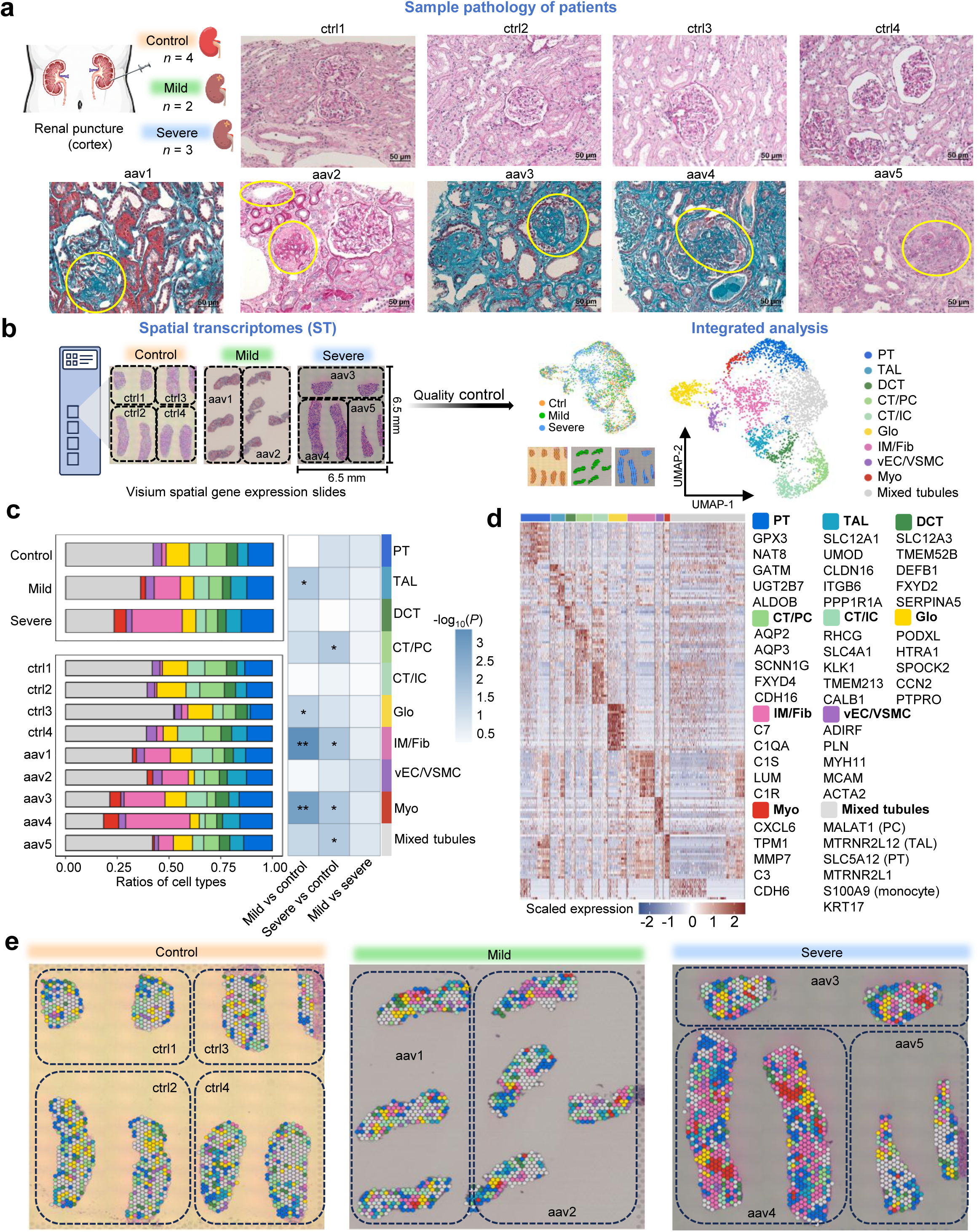
Overview of study design and clinical information in control (ctrl), mild and severe ANCA-associated vasculitis (AAV) datasets. (a) Histological staining of patient samples from control, mild AAV, and severe AAV groups. There were three groups of renal cortex tissues and corresponding histochemical staining images extracted from 4 (control), 2 (mild AAV) and 3 (severe AAV) patients respectively. Control patients are labeled as ‘ctrl’. The aav1 patient (mild group) exhibiting segmental glomerular sclerosis (Light microscopy, Masson trichrome-stained section). The aav2 patient (mild group) demonstrating glomerular global sclerosis with renal tubular atrophy (Light microscopy, PAS-stained section). The aav3-5 patients (severe group) showing crescentic formations in glomeruli (Light microscopy, aav3 & aav4: Masson trichrome; aav5: PAS). Yellow circles highlight pathological regions in all panels. Scale bars represent 50 μm. (b) Schematic workflow showing the study design and the subsequent spatial transcriptome analysis. The tissue sections were overlaid on Visium slides and sequenced. The data underwent quality control and subsequent downstream integrated analysis. UMAP plot showing the unsupervised clusters with cell type annotations assigned in integrated datasets. PT, proximal tubule; TAL, thick ascending limb; DCT, distal convoluted tubule; CT/PC, connecting tubule/principal cell; CT/IC, connecting tubule/intercalated cell; Glo, glomerulus; IM/Fib, immune cell and fibroblast; vEC/VSMC, vascular endothelial cell/vascular smooth muscle cell; Myo, myofibroblast. (c) Ratios of cell types in each group (left). The cell proportions in all patient samples were compared using Student’s *t*-test between the control, mild, and severe groups. The –log₁₀(*P*) are shown in the heatmap. * *P* ≤ 0.05, ** *P* ≤ 0.01. (d) Heatmap showing the top 20 most differentially expressed genes (DEGs) for each cell type in integrated datasets. For each cell type, five representative markers were highlighted. (e) H&E images overlayed by the annotated cell types in control, mild and severe AAV datasets.

We generated spatial transcriptomic (ST) profiles from kidney biopsies of control and AAV patients using the Visium platform, analysing 2-4 tissue sections per group (Fig. 1b). After standardizing quality control, we obtained 3,850 high-quality spots (1,413 in controls; 890 in mild AAV; 1,277 in severe AAV). Severe AAV samples exhibited increased transcript and gene counts per spot, accompanied by reduced mitochondrial content, consistent with altered composition and activation (Extended Data Fig. 1a).

Using unsupervised clustering and marker gene expression, we identified 10 distinct cell types from the integrated dataset (Extended Data Fig. 1e, Extended Data Fig. 2 and Supplementary Table 2), which were visualized in a UMAP plot and spatially mapped across kidney sections (Fig. 1b,e). These included proximal tubule (PT), thick ascending limb (TAL), distal convoluted tubule (DCT), connecting tubule/principal cells (CT/PC), connecting tubule/intercalated cells (CT/IC), glomerulus (Glo), immune cells/interstitial fibroblasts (IM/Fib), vascular endothelial/smooth muscle cells (vEC/VSMC), myofibroblasts (Myo), and mixed tubules. Cell type distributions were consistent across patients, and, as expected, controls showed fewer IM/Fib cells and almost no Myo cells (Extended Data Fig. 1b). The IM/Fib and Myo clusters showed the most pronounced changes among cell types in both mild and severe AAV patients compared to controls (Fig. 1c), with progressive expansion correlating with increasing disease severity and associated pathological features such as immune infiltration and fibrosis. The IM/Fib cluster was most abundant in the severe AAV group. In contrast, glomerular (Glo) spots were reduced in AAV samples, especially in the mild group, suggesting a potential link to early glomerular sclerosis. While TAL, CT/PC, and mixed tubule populations showed some variability across conditions, their changes were less prominent. Notably, the mixed tubule cluster co-expressed markers from multiple cell types (e.g., *MALAT1* for PC, *SLC5A12* for PT, *S100A9* for monocytes) (Fig. 1d), had the lowest transcript and gene counts, and exhibited high mitochondrial content (Extended Data Fig. 1d), were flagged as low-quality by pre-specified QC thresholds (genes/UMIs/% mito) and excluded a priori from downstream analyses.

We identified 2,865 upregulated differentially expressed genes (DEGs) across the 10 annotated kidney cell types (Supplementary Table 3). The top 20 DEGs showed distinct, cell-type-specific expression patterns (Fig. 1d, with strong concordance between our ST and scRNA-seq profiles from the Kidney Precision Medicine Project (KPMP) atlas^21^ (Supplementary Table 3). The IM/Fib and Myo clusters exhibited unique molecular signatures, including enrichment in complement pathway components (*C7, C1QA, C1S, C1R, C3*) and extracellular matrix organization (*LUM, MMP7*) (Extended Data Fig. 1f). Myo cells also showed upregulation of genes related to immune response (*CXCR6*), wound healing (*TPM1*), and cell junction assembly (*CDH6*). Our cell type annotations were further validated using scRNA-seq references from DISCO^62^ and the Kidney Cell Atlas (KCA) (https://www.kidneycellatlas.org/), confirming strong agreement with established marker gene expression (Extended Data Fig. 1c).

This ST profiling of AAV kidney biopsies revealed the severity-dependent expansion of immune-fibroblast and myofibroblast populations, as well as reduced glomerular signatures and distinct cell-type-specific gene expression programmes enriched for complement activation and fibrosis-related pathways.

### Delineating pathogenic AAV-responsive regions during disease progression

To characterise AAV-associated cell types *in situ*, we defined “disease-responsive” regions integrating changes in cell type proportions (Fig. 1c) with the expression of canonical AAV-related pathological marker genes (Supplementary Table 4 and Extended Data Fig. 3b). The AAV-responsive regions identify four key cell types – IM/Fib, Myo, Glo, and vEC/VSMC – as AAV-responsive (see Methods), which were consistently present across AAV patients, mirroring the distribution of injury markers (Extended Data Fig. 3b-d). These cell populations were spatially mapped across control, mild, and severe AAV groups (Fig. 2a), showing proximity within tissue sections and adjacent positioning in UMAP space (Extended Data Fig. 3a), which suggests shared transcriptional features. Consistent with our previous observations (Fig. 1c), IM/Fib and Myo populations expanded with increasing disease severity, while Glo proportions declined. Although vEC/VSMC cells were relatively sparse, they expressed immune-related genes (*C1S, C1R, C1QA–C1QC, IGHA1, JCHAIN*), injury markers (*IGFBP6, CST3*), and profibrotic genes (*COL1A2, COL3A1*), overlapping transcriptionally with both immune-fibrotic and glomerular compartments (Extended Data Fig. 3b). These four cell types, most strongly associated with AAV progression, were selected for focused downstream analyses.

**Fig. 2:**
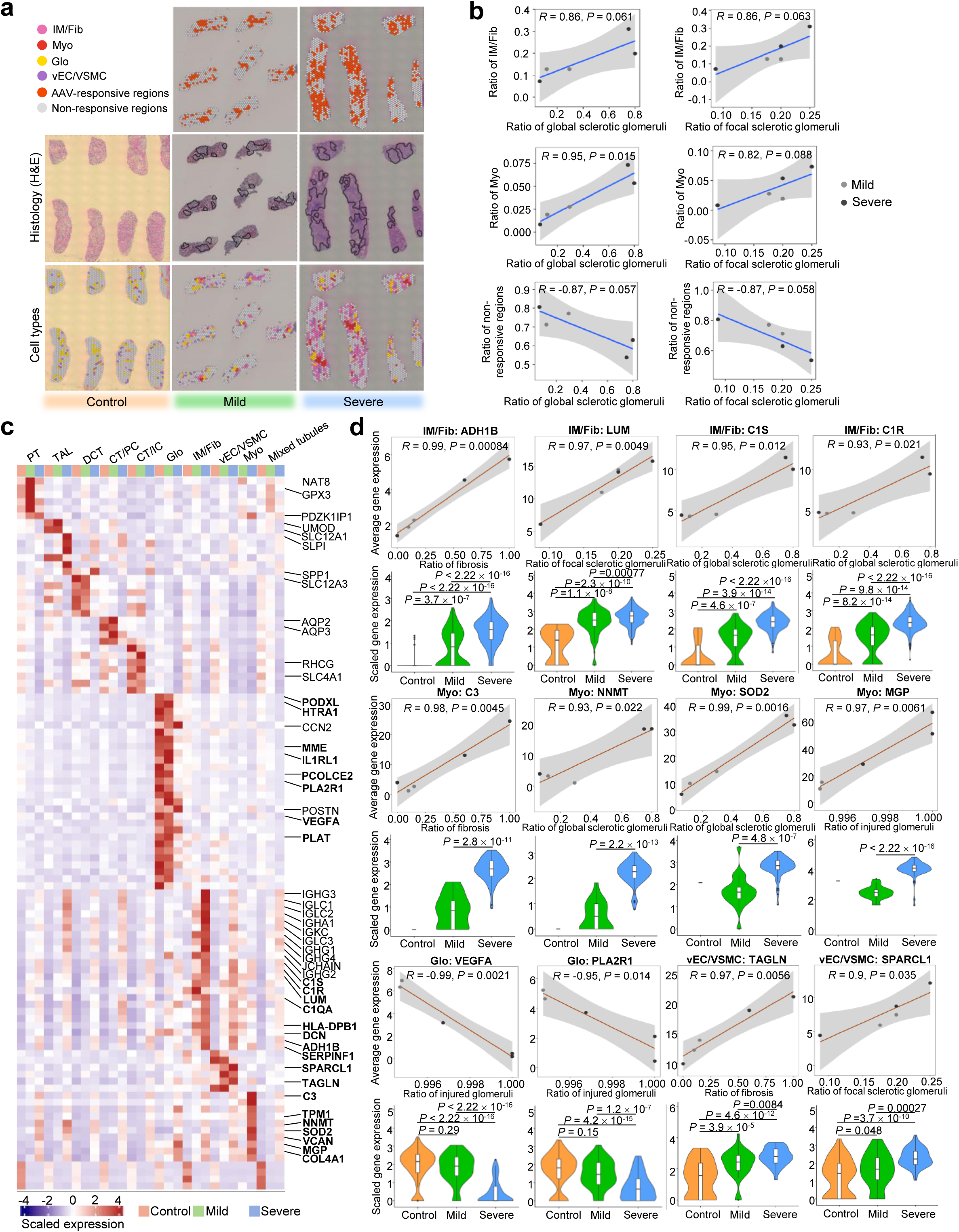
Identify and characterise AAV-responsive regions involved in disease progression. (a) Highlighted AAV-responsive regions, the matched histology and related cell types in AAV samples. Control samples are provided as a comparison. (b) The scatter plot illustrates the correlation between the pathological indicators of the kidney sections from AAV patients (x-axis) and the ratios of representative cell types/regions (y-axis) in five AAV patient samples. Non-responsive regions encompass cell types lacking AAV-responsive regions. (c) The heatmap displays the expression of cell type markers across control, mild, and severe AAV groups. Representative markers for each cell type are indicated, with those correlated to pathological indicators from AAV-responsive regions highlighted in bold (Pearson’s correlation: *P* ≤ 0.05. (d) Scatter plots show the relationship between kidney pathology indicators in AAV patients (x-axis) and the expression levels of key cell type markers in AAV-responsive regions (y-axis). Violin plots compare the expression levels of these markers across control, mild, and severe groups in each AAV-responsive region. *P*-values were determined using a Student’s *t*-test.

To assess whether AAV-responsive cell types are linked to disease pathology, we correlated cell type proportions from ST with histopathological indicators from diagnostic kidney biopsies (see Methods). Among all cell types, IM/Fib and Myo showed the strongest positive associations with key pathological features, including focal/global glomerulosclerosis, glomerular injury, and interstitial fibrosis (Extended Data Fig. 3e). IM/Fib proportions correlated strongly with both focal and global sclerosis, while Myo showed the highest correlation with global sclerosis (R = 0.95, p = 0.015) (Fig. 2b). In contrast, non-responsive regions exhibited negative correlations with these pathological measures. While PT cells also showed positive associations with injury, they were not prioritized due to minimal expression of AAV-relevant markers (Extended Data Fig. 3b). Glo and vEC/VSMC populations displayed distinct correlation patterns: Glo was positively associated with crescentic glomeruli, whereas vEC/VSMC showed inverse correlations. Together, these findings highlight dynamic shifts in IM/Fib and Myo populations, which may reflect their association with glomerular injury and tubulointerstitial fibrosis in AAV.

### Markers from AAV-responsive regions correlate with pathological indicators

We identified differentially expressed marker genes in AAV-responsive cell types (Glo, IM/Fib, Myo, and vEC/VSMC) that significantly correlate with pathological indicators (*R*² > 0.7, *P* < 0.05) (Fig. 2c,d and Supplementary Table 6). Tubular (e.g., *UMOD*, *SLC12A3*) and glomerular (e.g., *PODXL*, *HTRA1*) markers were markedly reduced in severe AAV, reflecting extensive loss of renal architecture. Notably, *VEGFA* and *PLA2R1*, podocyte-enriched genes^22^, showed strong inverse correlations with glomerular injury, highlighting podocyte damage in AAV. In IM/Fib populations, immunoglobulin genes and classical complement components (*C1QA*, *C1R*, *C1S*) were upregulated. ECM-related genes (*LUM*, *DCN*, *SERPINF1*, *ADH1B*) were also elevated, with *ADH1B* and *LUM* showing the strongest associations (*R*² > 0.97) with fibrosis and focal glomerulosclerosis, respectively. Markers in Myo and vEC/VSMC populations, including *VCAN*, *MGP*, *COL4A1*, *TPM1* and *C3*; *NNMT* and *SOD2*; as well as *SPARCL1* and *TAGLN,* were significantly upregulated in severe AAV and positively correlated with fibrotic and glomerular injury indices. *C3* had the highest correlation with fibrosis, while *MGP*, *NNMT,* and *SOD2* were linked with glomerular injury.

These cell types and marker associations with renal injury were independently confirmed in a separate AAV ST dataset^12^ (see Extended Data Fig. 3f–j), supporting their potential as diagnostic biomarkers for AAV in renal biopsies.

### Immune activation and extracellular matrix remodeling in AAV-responsive regions

We performed pairwise differential expression analysis across control, mild, and severe AAV groups in ten different kidney cell types. Disease-associated transcriptional changes were quantified by overlapping DEGs with cell-type-specific markers (Supplementary Table 2 and Fig. 3a). AAV-responsive regions showed a progressive increase in upregulated marker genes with disease severity, particularly in the IM/Fib compartment (severe *vs* control: 65%; severe *vs* mild: 48%). While most Glo markers were downregulated, a small subset was upregulated in severe AAV. Tubular markers were broadly downregulated, except for modest increases in PT and TAL-specific genes.

**Fig. 3:**
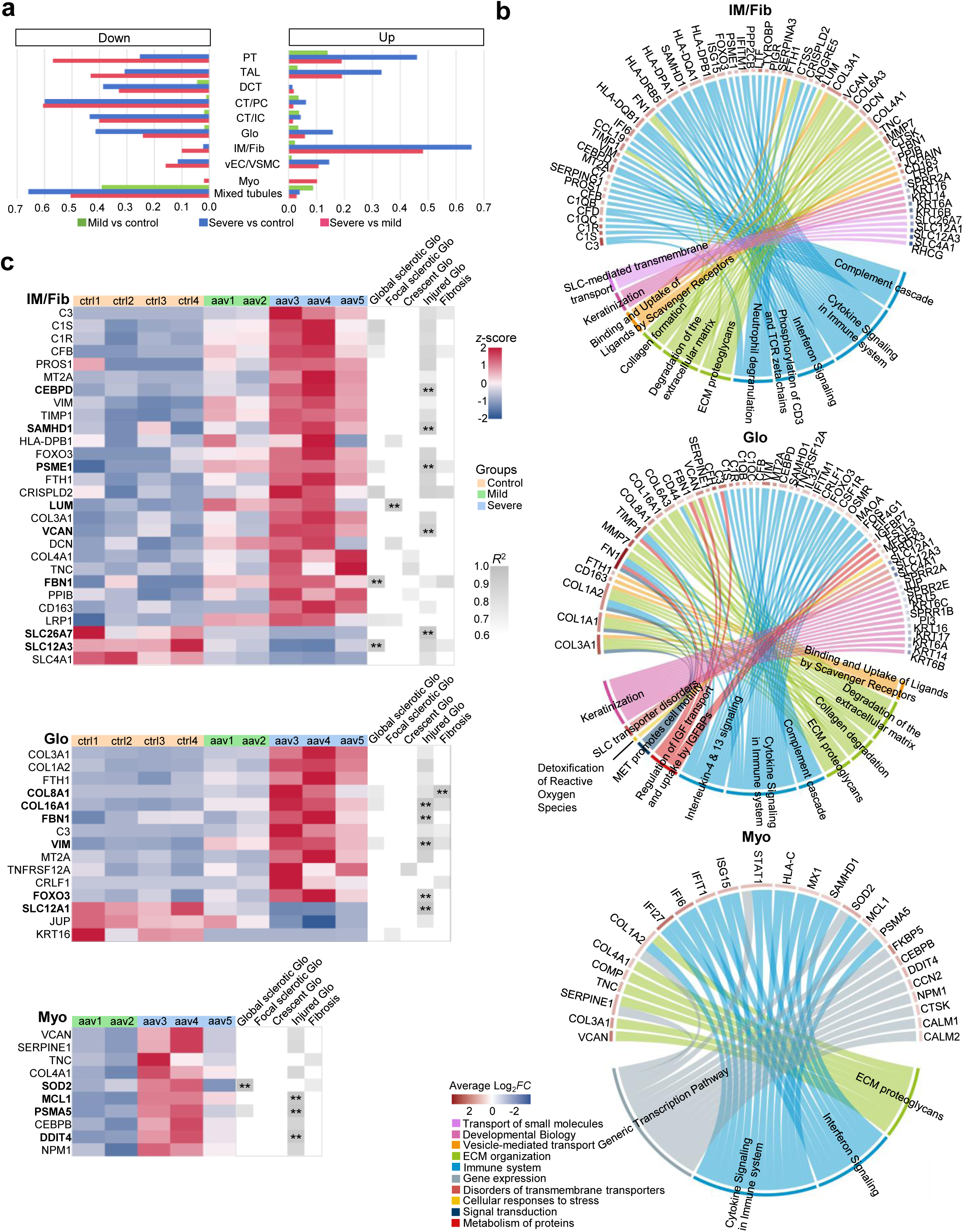
Functions of differentially expressed genes (DEGs) in disease-responsive regions and their relationships with clinical indicators. (a) The bar plot shows the ratios of up/down-regulated DEGs to cell type markers among control, mild, and severe groups. (b) Circular plots showing the top DEGs in AAV-responsive regions (IM/Fib, Myo and Glo) and their associated key biological functions, as identified through Gene Set Enrichment Analysis (GSEA) in the Reactome database. All functions are colour-coded on the outer ring to distinguish higher-level Reactome pathway categories, and the average log2 fold change (log_2_*FC*) for each gene is also annotated. (c) Heatmap (left) displaying the *z*-score of the expression levels of the top DEGs from AAV-responsive regions (IM/Fib, Myo and Glo) in each patient, which are correlated with clinical indicators (*R*² > 0.2, *P* < 0.05). The right heatmap illustrates the *R*² values representing the correlation between the DEGs and these clinical indicators. ** *P* < 0.01. The DEGs with the most significant correlation with indicators are highlighted in bold font.

To explore functional implications of these transcriptional changes, we performed Reactome-based^23^ gene set enrichment analysis (GSEA) on the DEGs from AAV-responsive regions (severe vs control; Fig. 3b and Extended Data Fig. 5a). We identified immune activation and ECM remodeling as the most significantly enriched pathways across all compartments. Core immune pathways, including complement activation, cytokine signaling, and interferon responses, were upregulated across the IM/Fib, Glo, Myo, and vEC/VSMC regions. However, the specific DEGs in these enriched pathways varied by cell type. Specifically, the IM/Fib region showed signatures of T cell activation and neutrophil degranulation, consistent with known immune infiltration in AAV^24,25^. In the Glo region, IL-4 and IL-13 signaling was enriched, suggesting a STAT6-mediated immune regulation and fibrosis, as previously reported^26^. ECM remodeling pathways, including collagen synthesis, proteoglycan deposition, and matrix degradation, were also broadly upregulated in all AAV-responsive regions (Fig. 3b and Extended Data Fig. 5a). A conserved set of ECM-related genes (*TIMP1, FN1, COL3A1, VCAN, COL6A3, COL4A1, TNC, MMP7, FBN1, COL1A2,* and *SERPINE1*) was identified across compartments, while others were region-specific (e.g., *LUM, DCN* in IM/Fib; *COL8A1, CD44* in Glo). Furthermore, the downregulated DEGs were primarily involved in development, transporter function, and protein metabolism, indicating impaired renal function. Correlating DEGs with pathology revealed that genes most strongly associated with clinical indicators (e.g., glomerular injury, fibrosis) were concentrated in IM/Fib, followed by vEC/VSMC, Glo, and Myo (Fig. 3c and Extended Data Fig. 5b). Notably, *LUM* (IM/Fib) correlated with focal glomerulosclerosis, while *FBN1* and *SLC12A3* (IM/Fib), *SOD2* (Myo), *C1R* (vEC/VSMC), and *COL8A1* (Glo) were most associated with global sclerosis and fibrosis.

Together, these findings suggest that immune activation and ECM remodeling are central to AAV pathogenesis, with both shared and region-specific molecular programmes across kidney compartments.

### Network analysis reveals coordinated immune and fibrotic pathways in AAV-responsive compartments

To identify coordinated transcriptional programmes in AAV-responsive regions, we used hdWGCNA on our ST data from severe AAV samples, which yielded 13 distinct gene co-expression modules (Extended Data Fig. 6a,b). Four of these (M2, M3, M8, M11) were enriched in AAV-responsive cell types: M2 and M11 in IM/Fib, M3 in Myo, and M8 in Glo (Fig. 4a). Among these, M11 was highly specific to IM/Fib, while M2 was also moderately active in Myo and Glo (Fig. 4b). M3 was mainly expressed in Myo, and M8 showed exclusive upregulation in Glo.

**Fig. 4:**
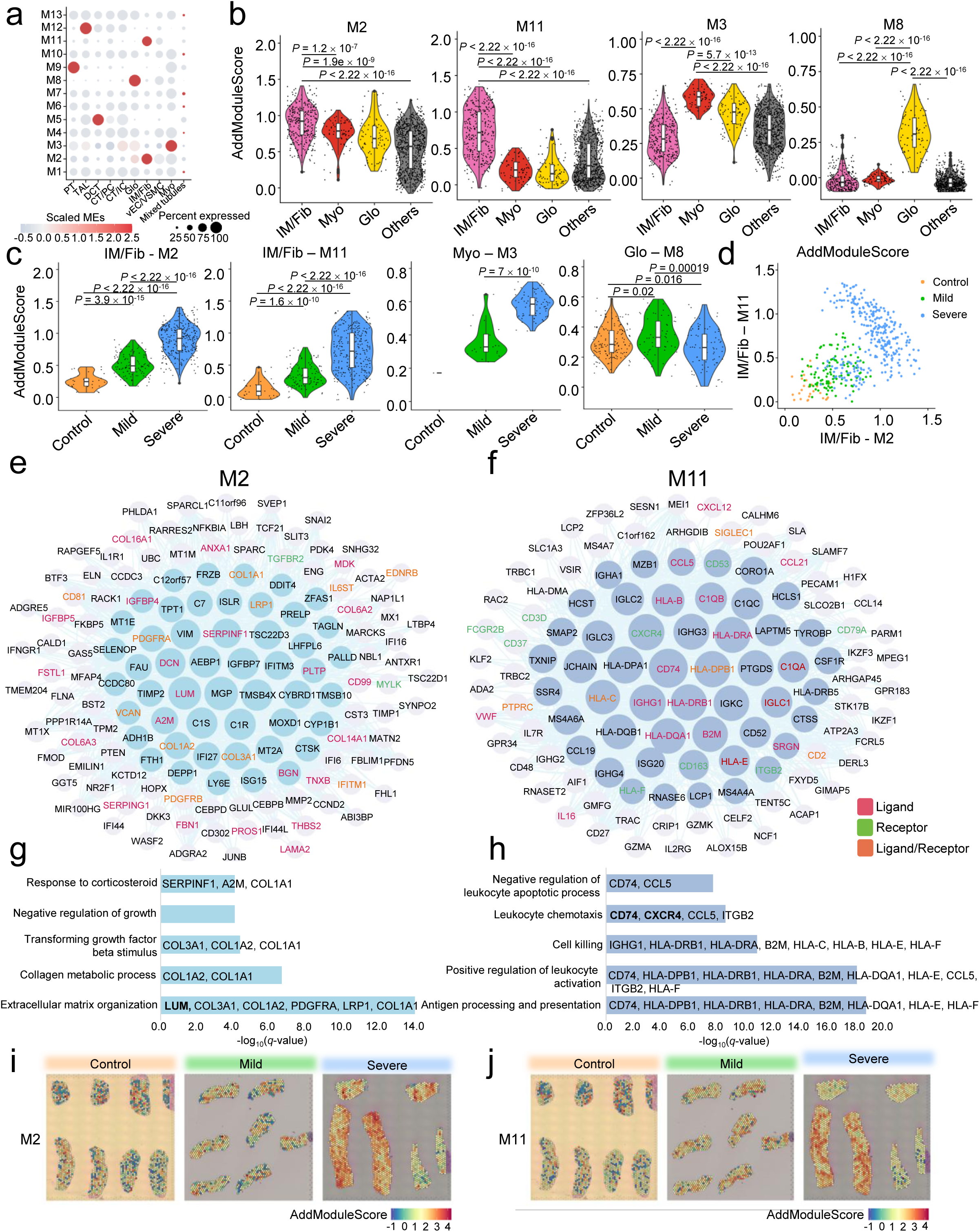
High-dimensional weighted gene co-expression network analysis (hdWGCNA) further characterises the gene co-expression networks in AAV-responsive regions of spatial transcriptomic data. (a) hdWGCNA identified 13 co-expression modules in the severe AAV group, with module eigengenes (MEs) representing the dominant expression patterns of each cell type. (b) Violin plots display expression of module features (top 100 genes) in AAV-responsive cell types of the severe group, with M2 and M11 predominantly localized in the IM/Fib region, M3 in the Myo region, and M8 in the Glo region. Feature expression levels were computed using the AddModuleScore. *P*-values were determined using a Student‘s *t*-test. (c) Violin plots show the comparison of module feature expression levels in each corresponding region across the control, mild and severe groups. *P*-values were determined using a Student‘s *t*-test. (d) Group-labeled scatter plot visualizing the correlation between M2 and M11 AddModuleScores in the IM/Fib region. (e-f) The network shows the eigengenes in M2 and M11. The 50 hub genes are represented by coloured circles, while ligands and receptors are also highlighted. (g-h) Top 5 enriched gene ontology biological process (GOBP) pathway from M2 and M11. Ligands or receptors from hub genes involved in each pathway are labeled in the figure. (i-j) Module activity patterns were spatially mapped using SpatialFeaturePlots, displaying the tissue distributions of M2 and M11 features generated by AddModuleScore.

We then assessed module activity using eigengenes and AddModuleScores derived from the top 100 hub-ranked genes (kME) (see Methods for details), and used this metric to evaluate each module’s association with AAV severity. Notably, M2, M3, and M11 activities increased with disease severity in their respective cell types, while M8 in Glo showed no such trend (Fig. 4c,d). In all IM/Fib spots, M2 features increased linearly with disease severity, while M11 features were induced in disease states (Fig. 4d).

Pathway enrichment and hub gene analysis highlighted M2 and M11 as key networks in IM/Fib (Fig. 4e–h). Compared to M3 and M8 (Extended Data Fig. 6c–f), M2 was enriched for ECM organization, featuring hub genes such as *COL1A1*, *COL1A2*, *COL3A1*, *LUM*, *PDGFRA*, and *LRP1*. These genes are involved in collagen biosynthesis and TGF-β signaling, both of which are known drivers of fibrosis in AAV nephropathy^27–29^. Additionally, the identified response to the corticosteroid pathway may be associated with corticosteroid treatment in AAV patients^30^. M11 was related to immune activation, including antigen presentation (*CD74*), leukocyte chemotaxis (*CXCR4*), cytotoxicity, and apoptosis (Fig. 4h). Cellular communication analysis suggests that *CD74* and *CXCR4* might cooperate in directing immune cell infiltration (Extended Data Fig. 7a–c).

In summary, although the spatial resolution did not allow for the complete separation of immune and fibrotic cells, M2 and M11 showed overlapping spatial features (Fig. 4d and i–j), suggesting that immune cell infiltration and fibrotic remodeling occur in adjacent or colocalized regions within AAV kidneys.

### CXCR4-CD74 crosstalk in the IM/Fib region mediates leukocyte chemotaxis in AAV

Integration of ligand–receptor and hdWGCNA analyses identified *CXCR4* and *CD74* as hub genes in module M11, associated with immune activation in the IM/Fib region (Fig. 4f). CD74 and CXCR4 form a receptor complex, and using NICHES for intercellular interaction analyses, we found that cell–cell communication in IM/Fib increased with AAV severity (Extended Data Fig. 7a), with the *CD74*–*CXCR4* axis showing one of the most pronounced activations in severe AAV (Extended Data Fig. 7b), and enriched for leukocyte chemotaxis, the top pathway in M11 (Fig. 4h and Extended Data Fig. 7c). While *CD74* was consistently upregulated across all AAV groups, *CXCR4* was selectively elevated in severe AAV (Fig. 5c). Spatial transcriptomics also revealed the co-expression or adjacency of *CXCR4* and *CD74* in AAV kidneys, particularly in severe disease (Fig. 5a,b and Extended Data Fig. 8a,b). In severe AAV, the co-expression of *CXCR4* and *CD74* was strongly positively correlated (Fig. 5d), suggesting disease-phase-dependent induction of this signaling axis, which may mediate leukocyte chemotaxis and subsequent immune activation.

**Fig. 5:**
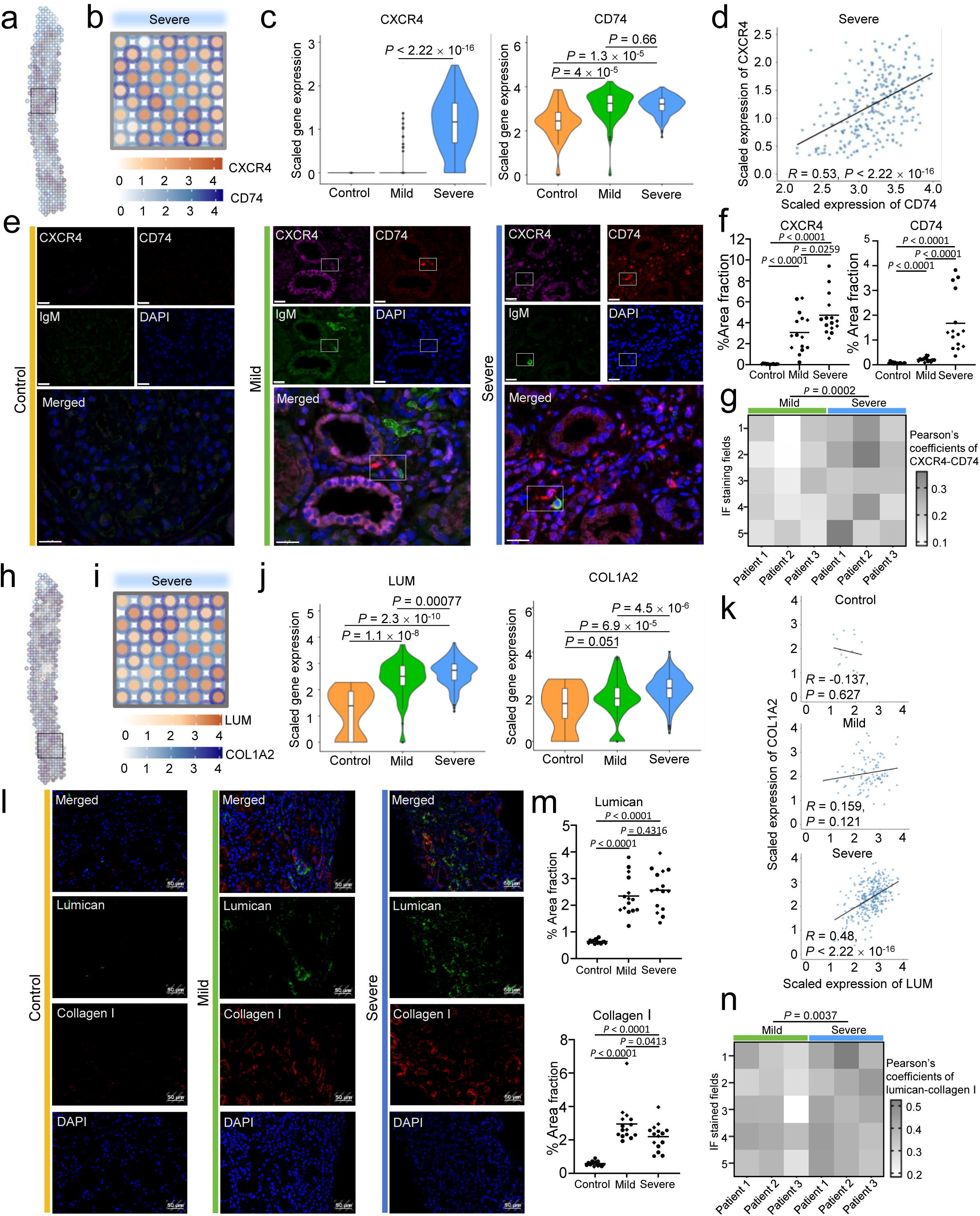
Ligand–receptor interactions reveal leukocyte chemotaxis and ECM remodeling in IM/Fib regions. (a) Spatially overlapped and scaled expression of *CXCR4* and *CD74* in a severe AAV patient. Expressions of *CD74* and *CXCR4* are visualized as an outer hollow circle and an inner solid circle, respectively. Fewer than 10 co-expressing spots were detected in mild AAV, precluding correlation analysis. (b) A close-up view (inset in a) reveals the spatial co-expression of *CXCR4* and *CD74* at spatial transcriptomic resolution. (c) Violin plots compare the expression levels of *CXCR4* and *CD74* across control, mild, and severe AAV groups in the IM/Fib. *P*-values were determined using a Student‘s *t*-test. (d) Scatter plots illustrate the correlation between *CD74* (x-axis) and *CXCR4* expression (y-axis) in spots co-expressing both markers (excluding spots where either *CD74* or *CXCR4* expression is zero) in the IM/Fib region of the severe AAV group. (e) Antibody-staining of CXCR4 (magenta), CD74 (red), and IgM (green) across control, mild, and severe groups. White boxes highlighted regions containing IgM⁺ cells (green) adjacent to CXCR4–CD74 co-localized cells (magenta/red) in representative mild and severe AAV patient samples. Scale bars represent 20 μm. (f) Relative protein expression levels of CXCR4 and CD74 (% area fraction of each antibody) across control, mild, and severe AAV groups. *P*-values were determined using a Student’s *t*-test. (g) Heatmap showing Pearson’s colocalization coefficients of CXCR4 and CD74 protein expression in mild and severe AAV groups, with *P*-values determined using Student’s *t*-test between all immunofluorescence (IF) staining fields of the two groups and labeled on the top of the heatmap. The Y-axis represents measurements from five IF fields per patient, while the X-axis indicates patients from each group (*n* = 3 per group). (h-k) Same as in (a–d), measured in *LUM* and *COL1A2.* (l) Antibody-staining of lumican (green) and collagen I (red) across control, mild, and severe groups. Scale bars represent 50 μm. (m-n) Same as in (f-g), measured in lumican and collagen I.

Immunofluorescence (IF) analysis corroborated these transcriptomic findings. CD74 and CXCR4 protein expression increased with disease severity, and colocalized near IgM-positive immune cells (Fig. 5e). *IGHM*, a marker of plasma/memory B cells exclusively expressed in AAV patients, was increased and positively correlated with *CXCR4* in severe AAV (Extended Data Fig. 8c–f). Spatial proximity analysis revealed that CD74/CXCR4 cells were 10–24 μm from IgM^+^ cells in the renal tubulointerstitium (Extended Data Fig. 8g,h), suggesting that CD74/CXCR4 may signal locally to attract nearby immune cells (a process consistent with paracrine chemotactic recruitment). We also reported AAV severity-dependent increases in CXCR4 protein expression and, to a lesser extent, in CD74 (Fig. 5f). The co-localization of CXCR4 and CD74, assessed by Pearson’s coefficients, was significantly higher in severe AAV (*P* = 0.0002; Fig. 5g), confirming a severity-linked interaction.

In summary, ST and IF analyses suggest that CXCR4–CD74 co-expression in IM/Fib regions is spatially linked to immune infiltrates and correlates with AAV severity, supporting their role in leukocyte chemotaxis and immune activation during disease progression.

### LUM–Collagen I/III co-localization correlates with AAV-associated renal fibrosis

In gene co-expression module M2, we identified *LUM* (lumican) as a central hub gene, with pathway and ligand-receptor analyses suggesting its interaction with *COL1A2* and *COL3A1* via *ITGB1*, implicating a role in ECM remodeling (Fig. 4e,g and Extended Data Fig. 7d). Spatial transcriptomics revealed strong spatial co-expression between *LUM* and both *COL1A2* and *COL3A1* within the IM/Fib region, either in the same or adjacent tissue spots (Fig. 5h, I and Extended Data Fig. 8i–l). Expression levels of *LUM*, *COL1A2*, and *COL3A1* increased progressively with AAV disease severity (Fig. 5j and Extended Data Fig. 8m), and all genes were significantly upregulated in AAV compared to controls (Fig. 5k and Extended Data Fig. 8n). These data suggest that LUM may contribute to pathological collagen deposition in AAV.

Immunofluorescence staining confirmed increased expression and co-localization of LUM with collagen I and III in AAV kidneys. Lumican and collagen I protein levels were elevated in both mild and severe AAV, with significantly stronger co-localization in severe cases (*P* = 0.0037, Fig. 5l-n). Similar patterns were observed for collagen III, which also showed enhanced co-localization with LUM in severe AAV (*P* < 0.0001; Extended Data Fig. 8o–q).

Together, these findings suggest that LUM is associated with fibrotic progression in AAV, potentially through spatial and functional interactions with fibrillar collagens involved in ECM accumulation.

## Discussion

Prior AAV studies have largely profiled circulating and infiltrating immune cells, with less attention to how disease programmes are organised within the renal microenvironment. Extensive immune infiltration in glomeruli has been reported^31^, and scRNA-seq analysis has revealed prominent populations of infiltrating B cells and plasma cells implicated in local immune responses within the kidney^32^. Further integration of ST and scRNA-seq has characterised inflammatory niches and highlighted pathogenic T cells in the kidneys of patients with ANCA-associated glomerulonephritis (GN)^12^. Building on these studies that identified inflammatory niches, we provide a pilot spatial map of AAV kidneys that localises immune–fibrotic activity *in situ,* and links it to histopathological severity at the patient level.

We defined four AAV-responsive renal regions, IM/Fib, Glo, Myo, and vEC/VSMC, and identified compartment-specific molecular signatures correlating with disease severity at the patient level. Among these, IM/Fib showed the most pronounced transcriptomic changes, including the highest number of differentially expressed genes, cell type markers, and strongest correlations with pathological indicators. The latter were evaluated at the patient level (to avoid pseudoreplication), i.e., for each cell type/region, we generated pseudo-bulk expression per patient and related these to histopathological indices using Spearman rank correlations with FDR control. Complement pathway activation was evident, with increased expression of classical pathway components *C1S* and *C1R* in IM/Fib and *C3* in Myo. While prior work emphasises alternative-pathway activation in AAV^33^, our increased classical components within IM/Fib are consistent with complement crosstalk in immune-complex-rich niches, potentially contributing to glomerular injury and thromboembolic risk in AAV^33–35^. We also identified ADH1B as a potential fibrosis-associated marker in IM/Fib, with minimal expression in controls and strong patient-level correlation with fibrosis scores. Given ADH1B’s roles in aldehyde/retinoid metabolism, we speculate that altered aldehyde handling could contribute to carbonyl stress and fibroblast activation and ECM deposition^36,37^ in AAV. This observation remains hypothesis-generating and warrants validation at the protein/activity level and dedicated functional analyses.

The immune response-associated module (M11) also includes immunoglobulins, HLA family members, and complement components that are enriched among cell type-specific DEGs in the IM/Fib region of the severe AAV group. In M11, the most prominent pathways play a central role in the development of kidney inflammation and renal injury in AAV. During leukocyte activation, ANCA binds to PR3 and MPO, leading to excessive neutrophil activation. These cells release large amounts of pro-inflammatory cytokines, damaging the vascular endothelium^7^. Activated monocytes and dendritic cells present antigens via MHC II (*CD74, HLA-DBP1, HLA-DRB1, HLA-DRA, HLA-DQA1*), activating T cells and establishing a persistent autoimmune response^6^. Vascular endothelial cells and immune cells also secrete chemokines (*CCL19, CCL5, CCL21*), attracting T cells and dendritic cells to accumulate at the sites of inflammation^38,39^. These processes collectively sustain tissue injury and loss of renal function. In addition, negative regulation of neutrophil and lymphocyte apoptosis may lead to sustained tissue damage (Supplementary Table 7).

CXCR4 and CD74 were co-expressed and co-localised within IM/Fib neighbourhoods. CXCR4 is a vital chemokine receptor implicated in leukocyte trafficking and fibrosis^40,41^. While prior studies support its involvement in kidney fibrosis, the potential interaction between CXCR4 and CD74 in AAV has not been reported. Rather than a classical ligand–receptor pair, CD74 and CXCR4 form a receptor complex that can be engaged by macrophage migration inhibitory factor (MIF)^42^ and tuned by CXCL12, promoting chemotaxis and inflammatory signaling^43^. Although MIF did not increase uniformly in our dataset, the spatial adjacency of CXCR4/CD74 with IgM⁺ cells and the severity-linked co-localisation are consistent with chemotactic recruitment. These features position the CXCR4–CD74 complex as a candidate tissue biomarker and a putative therapeutic axis in AAV, pending functional validation. We will quantify MIF/CXCL12 protein and test perturbations (e.g., blocking antibodies) to establish causality. However, further investigation is needed to identify the immune cell types involved and clarify whether these pathways also contribute to fibrogenesis.

In the ECM remodeling module (M2), key pathways including ECM organization, collagen metabolic processes, and TGF- β signaling were identified as drivers of fibrotic tissue injury^27,29,44^, which is consistent with findings from DEGs in the IM/Fib region of the severe AAV group. Beyond these major fibrosis-related pathways, we also identified pathways associated with the humoral immune response mediated by circulating immunoglobulin (*C1R, C1S, C7, SERPING1, CD81, SVEP1*) and the response to viruses (*IFITM3, IFI27, TPT1, ISG15, DDIT4*) within the M2 network (Supplementary Table 8). Interferon-stimulated genes (ISGs) were highly expressed in AAV, indicating abnormal activation of innate immunity and the maintenance of a pro-inflammatory and pro-fibrotic environment^45,46^. Hence, these findings suggest a crosstalk between fibrosis, innate immunity, and chronic inflammation within the AAV-responsive region.

A second key finding was the identification of *LUM* (lumican), a small leucine-rich proteoglycan and structural ECM component, as the hub gene in an ECM remodeling module (M2). *LUM* was strongly co-expressed with collagen genes *COL1A2* and *COL3A1* and spatially co-localized in fibrotic IM/Fib regions. These collagens are early markers of fibrosis^47^, and integrin-mediated interactions between LUM and collagen may contribute to excessive ECM deposition. Immunofluorescence confirmed LUM-collagen I/III co-localization and disease severity-linked expression. Interestingly, both LUM and collagen were upregulated even in mild AAV, suggesting early involvement in pathogenesis. Although LUM is not specific to AAV (being implicated in other kidney fibrotic diseases)^48–50^, its early and progressive upregulation in AAV suggests it as a potential biomarker for incipient fibrosis, when used in combination with other molecular indicators.

Moreover, these cell type-specific markers, including CXCR4, CD74, and LUM, may offer added value in enhancing diagnostic granularity and enabling patient stratification in renal biopsies from AAV patients. Unlike conventional histopathological assessments, these transcriptomic signatures showed correlations with disease severity and pinpointed specific pathological features, including immune infiltration and fibrosis. For instance, *LUM* and its co-localization with *COL1A2/COL3A1* may reflect fibrotic remodeling, while *CXCR4*–*CD74* co-expression could be indicative of localized immune activation. While preliminary, these findings suggest that integrating such molecular markers with histopathology may refine staging and guide therapies, pending validation in larger independent cohorts. We postulate that these markers constitute candidate tissue biomarkers with complementary roles: diagnostic (refining lesion classification in biopsies) and prognostic (risk stratification by immune activation and fibrotic burden). As the cost of spatial transcriptomics (ST)-based assays continues to decline, we envisage a composite spatial severity score (e.g., IM/Fib M11 and M2 module activity combined with marker gene load) to complement conventional histopathology. In addition to the IM/Fib region, we identified additional disease-associated markers in the Myo, Glo, and vEC/VSMC compartments, many of which correlated with pathological features, expanding the landscape of potential diagnostic markers.

Nevertheless, we acknowledge several limitations in our study. First, AAV rarity and the challenge of obtaining high-quality diagnostic biopsies resulted in a small cohort; accordingly, suggesting CXCR4, CD74, and LUM as candidate biomarkers remains preliminary. Validation in larger, independent cohorts will be necessary to establish their diagnostic and prognostic utility. Second, 10x Visium captures transcripts from multiple cells per spot, which limits single-cell resolution and introduces potential ambient RNA contamination and spot-mixing. We mitigated these by conducting patient-level analyses (i.e., pseudo-bulk per cell type/region with Spearman correlations, and implementing Benjamini–Hochberg FDR control), applying reference-guided cell-type enrichment analyses using several datasets (e.g., KPMP^21^, DISCO^62^ and KCA, and Giotto hypergeometric test), and confirming key findings by conducting blinded immunofluorescence quantification. Third, our cohort may not equally represent PR3- and MPO-ANCA subtypes or adult AAV cases, limiting generalizability. None of the patients had received treatment at the time of biopsy collection (hence minimizing this confounder), future studies should expand to larger, stratified cohorts based on serotype and age to replicate and extend these findings. Finally, finer cellular resolution will benefit from next-generation platforms such as Xenium, MERFISH, or seqFISH+^51–53^ and validation of main findings in larger, treatment-annotated cohorts is warranted.

In conclusion, we provide a foundational spatial map of AAV kidneys, which extends the previous immune cell subsets analysis^12,32^. These findings allowed us to generate testable hypotheses and a framework to refine biopsy classification and can help advance our understanding of AAV nephropathy. Future work integrating higher-resolution spatial transcriptomics with multi-omics approaches could further delineate cell-type-specific responses, enabling precise mapping of immune, fibrotic, and parenchymal cell dynamics in AAV nephropathy.

## Methods

### Ethical compliance

We have complied with all ethical regulations related to this study. The use of human kidney biopsy specimens in this study was approved by the Committee on Research Ethics of the Children’s Hospital of Nanjing Medical University. All experiments involving human samples were conducted in accordance with all relevant guidelines and regulations. All participants (or their legal guardians) provided written informed consent, and participation was entirely voluntary.

### Human kidney cortex biopsy samples used in spatial technologies and immunostaining

This study involved renal cortex biopsies from patients with ANCA-associated vasculitis (AAV), approved by the Independent Ethics Committee (IEC) of Children’s Hospital of Nanjing Medical University (Approval #202008089-1). Kidney samples from patients with AAV and healthy controls were analysed using spatial technologies and immunostaining (clinical details are provided in Supplementary Table 1). Healthy control tissues were obtained from non-diseased regions of patients with hematuria. In this study, cryopreserved kidney tissue sections were utilised for spatial transcriptomic analysis, whereas conventional FFPE (formalin-fixed paraffin-embedded) sections were prepared for fluorescence immunohistochemical staining.

### Histopathological Analysis of Renal Tissues

Renal tissue samples were fixed in 4% paraformaldehyde (PFA) at room temperature for 48 hours and then processed for histological examination. Serial sections were prepared at a thickness of 2 μm for Periodic Acid-Schiff (PAS) and Masson’s trichrome staining, all performed according to established protocols^54^. These kidney samples were systematically evaluated for structural abnormalities in the glomerular and tubular basement membranes and fibrotic changes, with subsequent Berden classification to stratify histological severity in ANCA-associated glomerulonephritis (details are provided in Supplementary Table 1).

### Construction and sequencing of spatial gene expression libraries

Spatial transcriptomic analysis was performed using the Visium HD Spatial Gene Expression platform (10x Genomics). Human kidney biopsy samples were prepared according to the Visium Spatial Protocols – Tissue Preparation Guide (10x Genomics, CG000240). Subsequent processing was conducted using the Visium Spatial Tissue Optimization Reagents Kit (10x Genomics, CG000238) and the Gene Expression Reagent Kit (10x Genomics, CG000239). Three Visium slides were sequenced, comprising control (n = 4, two sections each), mild (n = 2, with three and four sections), and severe (n = 3, two sections each). OCT-embedded tissue sections (10 µm) were methanol-fixed, hematoxylin and eosin (H&E)-stained, and imaged with a Leica DMi8 microscope (10x magnification) following the 10x Genomics protocols (CG000241). Following permeabilization for 12 minutes, mRNA was captured within the fiducial capture areas of the Visium slides. cDNA libraries were generated via second-strand synthesis, and sequencing was performed on a Novaseq X Plus system (Illumina).

### Spatial transcriptomics data processing, filtering, and gene annotation

Spatial transcriptomics data were processed using the Space Ranger pipeline (10x Genomics, v2.0.0) with the GRCh38 human reference genome (refdata-gex-GRCh38-2020-A) to generate raw unique molecular identifier (UMI) count matrices, mapped to 55 μm barcoded spatial spots.

Gene-spot matrices from Visium samples underwent individual quality assessment and preprocessing in R Studio (v4.3.2) with STutility (v1.0.0)^55^ and Seurat (v5.0.1)^56^. For each group (control, mild, and severe), raw counts, histological images, spatial coordinates, and scaling factors were loaded into STutility. Metadata (group, patient identifiers, and section numbers) were annotated with the “ManualAnnotation” function. Low-quality tissue spots were removed using the “subsetSTData” function. Metadata were transferred to Seurat objects, removing genes detected in <10% of spots. Mitochondrial content was calculated but not used for filtering due to biopsy size and possible mitochondrial expression changes in AAV lesions.

For each Seurat object, the gene-spot matrix was normalized using the “NormalizeData” function, and 2,000 highly variable features were identified via the “FindVariableFeatures” function. Integration anchors were then established using canonical correlation analysis (CCA) through the “SelectIntegrationFeatures” and “FindIntegrationAnchors” functions, enabling the integration of datasets for control, mild, and severe groups with the “IntegrateData” function. Then, we performed clustering analysis following Seurat’s standard workflow. The integrated data was scaled using the “ScaleData” function. Then, a combined principal component analysis (PCA) was performed using the “RunPCA” function, selecting the first nine principal components (PCs) based on variance inflection points identified through the visualization from the “ElbowPlot” function. These PCs served as input for subsequent UMAP dimensional reduction by the “RunUMAP” function and neighbourhood graph construction by the “FindNeighbors” function. Finally, we applied cluster detection at a resolution of 0.4 by the “FindClusters” function, which identified ten distinct transcriptional clusters within our dataset. The "DimPlot" function was used to display UMAP plots, utilising the "umap" reduction method and organising the data by either "cell types" or "groups," or splitting them as required.

### Cell type annotation

Cell types were assigned by identifying differentially expressed genes (DEGs) for each cluster versus all others using Seurat’s “FindAllMarkers” function (MAST method) with a non-parametric Wilcoxon rank sum test: log_2_ fold change (*FC*) ≥ 0.25, and adjusted *P* ≤ 0.05^57^. Clusters were annotated based on marker genes, cross-referenced with literature markers (Supplementary Table 2) and the Kidney Tissue Atlas^21^. Ten cell types were identified: proximal tubule (PT: *MIOX*, *ALDOB*), thick ascending limb (TAL: *SLC12A1*, *CLDN10*), distal convoluted tubule (DCT: *SLC12A3*), connecting tubule/principal cell (CT/PC: *AQP2*, *AQP3*), connecting tubule/intercalated cell (CT/IC: *FOXI1*, *ATP6V1B1*), glomerulus (Glo: *NPHS2*, *PTPRO*), immune cell types and interstitial fibroblasts (IM/Fib: *CD74*, *IGHG1*), vascular endothelial cells/smooth muscle cells (vEC/VSMC: *MYH11*, *ACTA2*), myofibroblasts (Myo: *COL8A1*), and mixed tubules. Cellular composition was calculated as the proportion of spots per cell type relative to total spots for inter-group comparison, and per patient relative to total spots for patient-level analysis.

### Gene co-expression network analysis

High-dimensional Weighted Gene Co-expression Network Analysis (hdWGCNA) was employed to find gene co-expression modules specific to the AAV-responsive cell types and associated with disease progression. Using the hdWGCNA package (v0.3.3) with a soft power threshold of 9^58^, we performed WGCNA on the AAV severe group (Extended Data Fig. 6a). The optimal soft power threshold is selected when the scale-free topology model-fit achieves its maximum value while the median connectivity attains its minimum to identify clusters with the strongest intra-group connections and the weakest inter-group connections. Module eigengenes (MEs) represent the first principal component (PC1) of the gene expression profiles within a co-expression module, serving as a summary of the module’s overall expression pattern. The average expression levels of the top 100 co-expressed genes (based on kME value) in each module were calculated using the “AddModuleScore” function in the Seurat package. The kME value (module membership measure) is a key metric that quantifies the strength of a gene’s connection to its assigned module. Based on cell type annotation, four modules (M2, M3, M8, and M11) were enriched in AAV-responsive cell types and exhibited gene expression changes correlated with disease severity. The topological overlap matrix (TOM) of these modules was computed using the “GetTOM” function in hdWGCNA. Hub gene networks for each module were visualized using Cytoscape (v3.9.1), and ligand-receptor genes within each module were annotated based on human ligand-receptor pairs from the FANTOM5 database^59^.

### Differential expression analysis

Differential expression analysis was performed in Seurat using “FindMarkers” (MAST method) for each cell type between mild vs. control, severe vs. control, and severe vs. mild groups^57^. MAST (Model-based Analysis of Single-cell Transcriptomics) is a statistical framework that utilises generalized linear models (GLMs) to analyse two gene expression features: a binomial component for gene detection probability and a continuous component for transcript abundance. In the “FindMarkers” function, MAST improves differential expression analysis by integrating covariates with both gene detection rates and expression levels, hence increasing biomarker identification by accounting for technical uncertainty and cellular heterogeneity. DEGs were defined as log_2_*FC* ≥ 0.25, and adjusted *P* ≤ 0.05 (Wilcoxon rank sum test).

### Validation of cell type annotations

To validate cell type annotations, we performed hypergeometric test-based enrichment analysis using the Giotto package (v3.3.0)^60^. For each group (control, mild, and severe), Giotto objects were generated with Seurat metadata. References (Supplementary Table 9) were restricted to scRNA-seq data from healthy kidneys and cortex-derived cell types to avoid disease-related bias. Cell-type-specific marker genes were identified using Seurat’s “FindAllMarkers” function (MAST method) from the Kidney Cell Atlas mature kidney dataset^61^ and healthy kidney data in the Deeply Integrated Single-Cell Omics (DISCO) database^62^. Marker genes cutoff: log_2_*FC* ≥ 0.25 and adjusted *P* ≤ 0.05 (Wilcoxon rank sum test). We performed hypergeometric testing using Giotto’s "runHyperGeometricEnrich" function to generate enrichment scores, defined as -log_10_(*P*) per spatial spot. Mean scores for each annotated cell type were visualized as a heatmap.

### Identification of AAV-responsive regions

AAV-responsive regions were defined by two criteria: (1) cell type proportions differed significantly (*P* ≤ 0.05) between mild vs. control groups or severe vs. control groups; (2) AAV marker genes (Supplementary Table 4) were expressed in disease but not controls, with scaled average expression >0 in both mild and severe groups (Supplementary Table 5 and Extended Data Fig. 3b).

Although vEC/VSMC showed no significant proportion changes and had few spots, they were included in the AAV-responsive region because these cell types are associated with vasculitis according to the disease characteristics. Besides, mixed tubules were excluded because the spatial transcriptomics resolution/quality was insufficient to distinguish tubular subtypes. In summary, we defined IM/Fib, Myo, Glo, and vEC/VSMC as AAV-responsive regions, which meet the criteria above.

### Association between cell type and pathological indicators

The association between cell type proportions from integrated spatial transcriptomics and pathological indicators from patient biopsies was assessed using Pearson correlation using the ‘cor’ function from the stats package (v4.3.2), with significance determined via *P*-values within 95% confidence intervals from the ‘cor.mtest’ function in the corrplot package (v0.95). The pathological indicators were derived from quantitative metrics obtained from diagnostic biopsies (Supplementary Table 1). Results were visualized as a clustered heatmap by the ‘pheatmap’ function, with rows as pathological indicators, columns as cell types, and colour intensity indicating the Pearson correlation coefficient (Extended Data Fig. 3e). Statistically significances were annotated with asterisks (\**P* ≤ 0.1, ** *P* ≤ 0.05) and hierarchical clustering to identify cell types potentially linked to clinical features.

### Correlation between cell type markers/DEGs and pathological indicators

To evaluate correlations between cell-type-specific markers/DEGs derived from integrated spatial transcriptomics and pathological indicators from patient biopsies, we employed linear regression analyses using the ’lm’ function from the stats package implemented in a computational loop. For each gene-pathology indicator pair, separate regression models were fit, with *R*² and *P*-values. Correlations meeting dual thresholds (*P* < 0.05 and *R*² > 0.2) were considered biologically meaningful.

### Functional enrichment analysis

To determine the key biological pathways associated with each cell type, we performed over-representation analysis (ORA) on the commonly upregulated DEGs from each cell type using the clusterProfiler package (v4.8.3)^63^. Enrichment was performed against Gene Ontology Biological Process (GOBP) terms^64^, with redundant GO entries removed using the simplify function (similarity cutoff=0.6). Pathways with *q*-value ≤ 0.05 were considered significant. WGCNA modules and ligand–receptor functions in AAV-responsive regions were analysed with the same method. To investigate the functions of DEGs in the AAV-responsive regions, we performed gene set enrichment analysis (GSEA) on the DEGs^63,65^, which were ranked by their average Log_2_*FC* values from each AAV-responsive cell type. This analysis was performed using the “gesPathway” function from the ReactomePA package (v1.44.0)^66^. The normalized enrichment scores (NES) were weighted based on the overlap between the input gene sets and Reactome Pathway’s canonical functional gene sets. Multiple testing correction was applied using the Benjamini-Hochberg (BH) method, with significance thresholds set at an FDR ≤ 20% and a minimum gene set size (minGSSize) of 10. The most significant DEGs and associated functions were visualized using the circlize package (v0.4.16).

### Human AAV spatial transcriptomics reference dataset used to validate key features of AAV-responsive regions

A recently available human AAV spatial transcriptomics reference confirmed the characteristics of AAV-responsive regions in our dataset^12^. The reference datasets comprised 10 predefined normal renal cell types (including CT/PC, CT/PC/IC, DCT/CT, Glo, LOH, PT, PT/DCT, PT/TAL, tubulointerstitium, and vasculature) and 2 AAV-responsive cell populations (inflamed Glo and tubulointerstitium), derived from 19 ANCA-crescentic glomerulonephritis (GN) patients and 8 controls (Supplementary Table 9). We evaluated the similarity between our spatial transcriptomics data and the reference’s gene expression profiles using the fgsea package (v1.26.0). The DEGs identified by the “FindMarkers” function in the reference dataset (between controls and ANCA-GN patients, with log_2_*FC* ≥1 and adjusted p ≤ 0.05) and the up-regulated DEGs from our AAV severe group (ranked by log_2_*FC*) were used as input for analysis. The NES was then calculated to show the enrichment of cell-type features from the reference dataset within our cell types. The key cell type markers identified in our AAV-responsive regions were further validated within the spatial reference dataset.

### Cell-cell communication analysis using NICHES

Ligand-receptor (L-R) cell-cell communication analyses were performed using the NICHES package (v1.0.0)^67^. NICHES analyses the gene expression matrix with spot metadata from spatial transcriptomics as input and generates interaction matrices as output using the “RunNICHES” function. In the matrices, each row represents a known ligand-receptor pair from OmniPath databases^68^, while each column shows potential cell-cell communication events. The interaction strength for each pair is calculated by multiplying the ligand’s expression level in the sending cell by the receptor’s expression level in the receiving cell. The ligand-receptor interaction matrices can be analysed in Seurat using the “FindMarkers” function, allowing for a comparison of cell-type-specific ligand-receptor interactions between the AAV and control groups. The significance thresholds were set using the non-parametric Wilcoxon rank sum test with the following cutoffs: log_2_*FC* ≥ 0.25 and adjusted *P* ≤ 0.05.

We used the InterCellar package (v2.6.0) to construct interaction networks, visualizing cell-cell communication patterns^69^. In these networks, nodes represent distinct cell types from each experimental group, while edges indicate either autocrine (within the same cell type) or paracrine (between different cell types) interaction frequencies.

### Immunofluorescence staining

Immunofluorescence (IF) analysis was conducted on 3-μm paraffin-embedded kidney sections. Tissue samples were derived from three groups: controls, mild, and severe AAV patients (n = 3 in each experimental group). The IF staining for triple-labeling tissue sections using a TSA-based four-colour kit system (Abclonal, #RK05903) begins with deparaffinization by baking slides at 65°C for 30 minutes, followed by sequential immersion in xylene (37°C), absolute ethanol, and an ethanol gradient (95%, 85%, 70%) for rehydration, then rinsed in PBS. Antigen retrieval is performed in Tris-EDTA buffer (pH 9.0, Beyotime, #P0084) using an antigen retrieval cooker (Aptum, 2100 Retriever) for 20 minutes, cooled to room temperature (RT), and washed with PBS. Endogenous peroxidase activity was quenched with 3% H_2_O_2_ (ORIGENE, #PV-9000) for 20 minutes at RT, followed by PBS rinses and blocking with 5% BSA (Beyotime, #P0102) for 1 hour at RT. Primary antibodies are applied sequentially over three days: Day 1, CXCR4 antibody (Abcam, #ab124824, 1:100); Day 2, CD74 antibody (Invitrogen, #14-0747-82, 1:100); Day 3, IgM antibody (ORIGENE, #24052901, 1:30, direct green fluorescence without secondary antibody). First antibodies (without IgM) are incubated overnight at 4°C, then washed, and treated with HRP-conjugated secondary antibody (Abclonal, #RK05903-5) for 50 minutes at RT. IF staining used stains TYR-570 for CD74 antibody (red, Abclonal, #RK05903-2); TYR-690 for CXCR4 antibody (violet, Abclonal, #RK05903-3), amplified with TSA+enhancer (1:200, Abclonal, #RK05903-4). After each round, the antibodies are stripped using a 95°C EDTA buffer for 40 minutes, and the cycle (peroxidase blocking to TSA) is repeated for subsequent targets. Finally, these sections are mounted using an antifade mounting medium with DAPI (Vector Laboratories, #H-1200). Slide images were acquired using a Panoramic MIDI slide scanner (3DHISTECH, #PMIDI23C3001) fitted with a Grasshopper3 USB 3.0 camera (FLIR, #GS3-U3-51S5M-C) and a Zeiss 20X Plan-Aprochromat objective. Image capture parameters included an 8-bit depth and variable exposure times ranging from 8 to 60 ms. Post-acquisition processing was performed using CaseViewer software (3DHISTECH).

The IF staining for double-labeling tissue sections using a TSA-based three-colour kit system (Abclonal, #RK05902). The procedure for IF staining was performed following the same protocol as described above. Primary antibodies are applied sequentially over two days: Day 1, the lumican antibody (Proteintech, #10677-1-AP, 1:400) or the CXCR4 antibody; Day 2, the collagen I and collagen III antibodies (Bioss, #BS-10423R and #BS-0549, 1:400) or the CD74 antibody. IF staining used stains TYR-520 for CXCR4 and lumican antibodies (green, Abclonal, #RK05903-1); TYR-570 for CD74, collagen I, and collagen III antibodies (red, Abclonal, #RK05903-2). Fluorescence was detected using an X-cite Series 120 Q unit (Excelitas, USA) attached to a light microscope Imager A2 (Zeiss, Germany) with an Axiocam 503 colour digital camera (Zeiss, Germany). Five randomly selected fields per section (200X magnification) were imaged, with all acquisitions performed under standardized optical and exposure conditions. Quantitative analysis was conducted using ImageJ (v1.54k, National Institutes of Health, USA)^70^.

### Statistical analysis

Statistical analyses were performed using R Studio (v4.3.2) or GraphPad Prism software (v10.4.1; GraphPad Software, LLC, www.graphpad.com). Continuous variables are expressed as mean ± standard deviation (SD), with between-group differences evaluated by two-tailed, unpaired Student’s *t*-tests. Statistical significance was defined as a *P* < 0.05, unless otherwise indicated. Differences in cell type proportions were assessed via Student’s *t*-test, with significance levels denoted as * (*P* ≤ 0.05) and ** (*P* ≤ 0.01).

### Packages used to generate spatial transcriptomics figures

The top DEGs for each cell type were plotted as a heatmap using the "DoHeatmap" function in Seurat, representing the integrated datasets. Other heatmaps were generated using the pheatmap package (v1.0.12), employing the “pheatmap” function with optimized parameters.

The spatial distribution of cell types or AAV-responsive regions in control, mild, and severe datasets was visualized using the “SpatialDimPlot” function in Seurat. The “SpatialFeaturePlot” function was used to reveal the scaled expression and spatial patterns of key cell type markers in AAV-responsive regions. Violin plots were created using the “VlnPlot” function in Seurat. All dot plots were generated using the “DotPlot” function in Seurat to display feature expression changes across different cell types.

Figures not mentioned before (such as bar plots, combined violin-box plots, and scatter plots) were generated using the ggplot2 package (v3.5.1) with optimized parameters.

## Data availability

The spatial transcriptomic data supporting the findings of this study have been deposited in the CNSA database and will be made publicly available upon publication.

## Code availability

All analyses were conducted using open-source software and packages, as cited in the Methods section. Additional information can be provided upon request.

## Supporting information

Extend1

