## Supplementary material for "Spatial transcriptomics reveals injury-responsive compartments and coordinated immune–fibrotic signaling in ANCA-associated renal vasculitis": Extend1

### Extended Data Fig.1

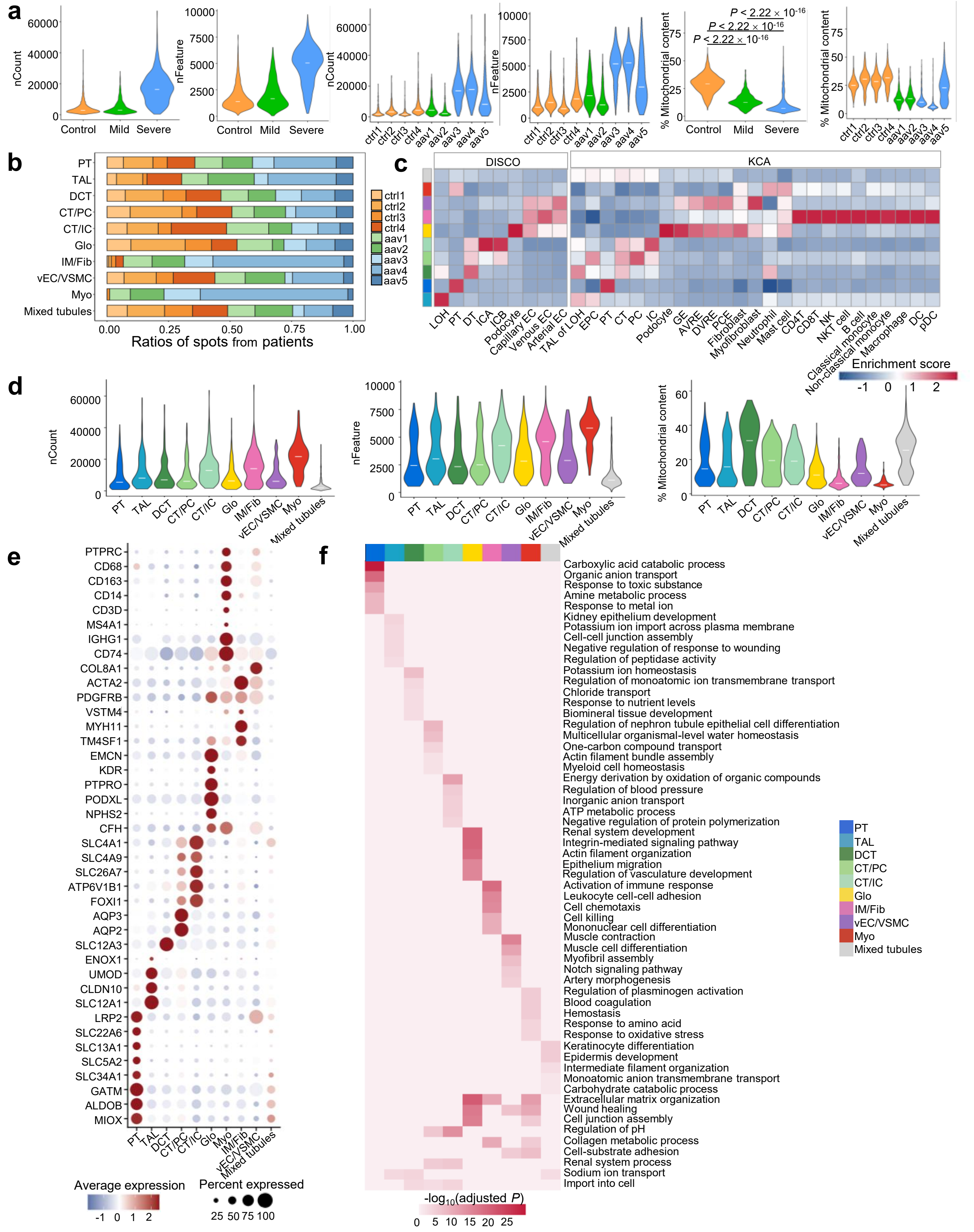


**Extended Data Fig.1 Quality control, annotation, and cell type validation in the integrated data.** (a) The QC metrics across groups and patients were shown in the violin plot (Wilcox test). (b) The ratio distribution of sequenced spots from each patient across cell types. (c) Giotto hypergeometric tests with two human kidney scRNA-seq reference datasets from the Deeply Integrated human Single-Cell Omics (DISCO) and Kidney Cell Atlas (KCA) databases across all clusters in the integrated datasets. The enrichment score is calculated based on the *P*-value from the hypergeometric test, $-\log_{10}(P)$, and finally applying *z*-score normalization. LOH, Loop of Henle; PT, proximal tubule; DT, distal tubule; ICA, intercalated cell type A; ICB, intercalated cell type B; EC, endothelial cell; TAL, thick ascending limb; EPC, epithelial progenitor cell; CT, connecting tubule; PC, principal cell; IC, intercalated cell; GE, glomerular endothelium; AVRE; ascending vasa recta endothelium; DVRE, descending vasa recta endothelium; PEC, peritubular capillary endothelium; DC, dendritic cell; pDC, plasmacytoid dendritic cell. (d) The QC metrics across cell types from the integrated dataset were shown in the violin plot. (e) Dot plot representing gene expression of canonical cell type markers in integrated datasets. (f) Heatmap of top 5 GOBP pathways for each cell type and top shared GOBP pathways among cell types in integrated datasets by GO over-representation analysis.
